# Patterns in protein flexibility: a comparison of NMR “ensembles”, MD trajectories and crystallographic B-factors: Patterns in protein flexibility

**DOI:** 10.1101/240655

**Authors:** Anthony Riga, Jasmin Rivera, David A. Snyder

## Abstract

Proteins are molecular machines requiring flexibility to function. Crystallographic B-factors and Molecular Dynamics (MD) simulations both provide insights into protein flexibility on an atomic scale. Nuclear Magnetic Resonance (NMR) lacks a universally accepted analog of the B-factor, however, a lack of convergence in atomic coordinates in an NMR-based structure calculation also suggests atomic mobility. This paper describes a pattern in the coordinate uncertainties of backbone heavy atoms in NMR-derived structural “ensembles” first noted in the development of FindCore2 (previously called Expanded FindCore: DA Snyder, J Grullon, YJ Huang, R Tejero, GT Montelione, *Proteins: Structure, Function, and Bioinformatics* 82 (S2), 219–230) and demonstrates that this pattern exists in coordinate variances across MD trajectories but not in crystallographic B-factors. This either suggests that MD trajectories and NMR “ensembles” capture motional behavior of peptide bond units not captured by B-factors or indicates a deficiency common to force fields used in both NMR and MD calculations. Additionally, a comparison of Cα B-factors with Cα coordinate variability in NMR “ensembles” and MD trajectories shows that NMR-derived coordinate uncertainties measure variability in atomic positions as well as crystallographic B-factors and superimpositions of MD trajectories do.

## Introduction

Large molecules and biomolecules can have a high degree of motional flexibility, affecting their function [1]. Common sources of information about protein flexibility and modes of motion include crystallographic B-factors [2], molecular dynamics (MD) simulations [3] and Nuclear Magnetic Resonance (NMR) spectroscopy, including relaxation measurements [4–7] and even chemical shift data [8,9].

Each of the above techniques for evaluating protein flexibility yields an incomplete picture of protein dynamics in solution. Crystallographic B-factors are affected by packing and other special features of the crystalline state [10]. In addition, many factors may reduce the intensities of the “reflections” in a protein crystal’s X-ray diffraction pattern, and hence elevated crystallographic B-factors may not only indicate macromolecular flexibility [11]. The quality of MD simulations is dependent on the quality of the seed structure and force field used, although recent efforts have applied MD simulations to NMR-derived structures [12], and NMR relaxation experiments provide a critical source of data for evaluating individual MD trajectories as well as the force fields and other methodological details of MD simulations [13,14]. The combination of multiple assessments of protein flexibility has proven particularly illuminating [15]: for example, the combination of NMR-relaxation data with MD simulations yields a detailed picture of protein dynamics and motional modes [16].

While lacking a universally accepted analog of the B-factor, the NMR based structure determination process itself nonetheless provides insight into protein flexibility. Atoms in loop residues and other flexible regions of a protein typically have fewer long-range “contacts” to atoms in other residues. This paucity of contacts leads to both increased flexibility of loop regions [17,18] as well as poor convergence for loop residue positions in NMR-based structure calculations [19–22], provided the structure refinement process does not lead to inaccurate rather than imprecise coordinates [23]. Moreover, the primary source of structural restraints in NMR-based structure calculations are NOESY experiments and fast motions reduce NOEs while intermediate time scale motion causes line broadening that can interfere with the identification of NOESY cross-peaks. Thus, NMR yields a paucity of restraints for particularly flexible regions of a protein leading to poor convergence in NMR-based structure determination, and coordinate uncertainties in an NMR-derived “ensemble” of structures, which strictly speaking measure the local reproducibility of the NMR-based structure determination process, reflect region-specific and perhaps even atom specific levels of flexibility [24].

While they can provide key insights into protein flexibility and dynamics, evaluation of uncertainties in protein structure coordinates inferred from NMR data is a non-trivial process. Typically, NMR-based structure calculations generate multiple (typically 10–40) models [22]. Such collections of structural models are called “ensembles”: although NMR ensemble generation can effect Boltzmann sampling [19–21], generally NMR ensembles, including those analyzed in this study, are *not* actually Boltzmann ensembles.

Calculation of coordinate variances requires the superimposition of NMR ensembles. However, inclusion of poorly converged coordinates can bias the superimposition process, reducing the applicability of the resulting coordinate variances [25,26]. Limiting the calculation of an optimal superimposition to a core atom set, determined in a superimposition independent manner using either circular variances of backbone dihedral angles [27] or an interatomic variance matrix [26,28], ensures calculation of optimal superimpositions and hence of appropriate coordinate uncertainties. Alternatively, assumptions concerning the distribution of coordinate variances can lead to model based superimposition methods such as THESEUS which assumes a multivariate Gaussian distribution of coordinate uncertainties [29,30].

Identification of a core atom set is a critical step in solving two distinct, albeit related, problems. Not only is identification of a core atom set an important step in calculating coordinate uncertainties via superimposition, but such core atom or residue sets also convey in which regions the NMR-based structure calculation process has converged [22]. As these two problems are different, their optimal solutions may differ slightly. For example, application of the FindCore method, which identifies core atom sets for use in assessing the *precision* of NMR ensembles, to the distinct, albeit related problem of identifying *well-converged* core atom sets for CASP10 [31,32] required extension of the FindCore method into an approach known as Expanded FindCore [22].

Software used in the CASP10 competition also required any residue with core atoms to have all backbone heavy atoms in the core. The process of modifying Expanded FindCore to meet this requirement revealed carbonyl oxygens from otherwise well-defined residues whose positions were poorly defined in NMR-based structure calculations. Given the relation between coordinate uncertainties in NMR-derived structures and physical flexibility as described above, this discovery raised questions about the high uncertainties (relative to other backbone heavy atoms in the same residue) of those carbonyl oxygens: how common are these relatively uncertain carbonyl oxygens and is this high relative uncertainty an artifact of the NMR-based structure determination process or is it indicative of a pattern in backbone atom flexibilities?

Addressing these questions requires comparison of NMR ensembles with complementary structural information, such as that obtained from crystallographic data, as well as with MD trajectories that provide insight into protein flexibility. Protein structures obtained by the North East Structure Genomics (NESG; http://www.nesg.org/) consortium facilitated this analysis: the NESG performed crystallization and HSQC screening in parallel for robustly expressed protein targets. This has resulted in over 40 NMR/X-ray crystal structure pairs [33,34]. Four of those pairs are subjected to further analysis here. Statistical analyses of backbone heavy atom coordinate variances observed in MD trajectories, seeded by crystal structures and superimposed using THESEUS [29,30], identifies the same pattern in relative coordinate uncertainties as observed in NMR ensembles superimposed using FindCore [26].

The analysis presented here demonstrates the persistence of a pattern in coordinate variances across structural “ensembles” obtained using multiple force fields (AMBER [35,36] and OPLS [37] in MD simulations, CNS [38,39] in NMR structure refinement), superimposition techniques and sampling schemes (restrained simulated annealing and similar schemes in NMR structure refinement as distinct from the unrestrained constant temperature approach used in MD). This persistent pattern does not (necessarily) occur in Crystallographic B-factors of backbone heavy atoms as demonstrated by its absence in the four crystal structures analyzed in this study. Even in the four proteins explored here, that the relatively high uncertainty of carbonyl oxygens (and also Cα atoms) persists, in almost all MD trajectories simulated in this study, indicates that the relatively high uncertainty of carbonyl oxygens (and also Cα atoms) is not solely an artifact of NMR-based structure determination. The pattern in backbone heavy atom coordinate uncertainties reflects either a physical reality of peptide bond motion not evident in crystallographic data or a shortcoming common to multiple force fields. If the latter explanation is true, the analysis presented here underscores that further improvements in force field parameterization are necessary for better prediction and calculation of protein structure and dynamics.

## Materials and methods

MD simulations were initiated using crystallographic structures retrieved from the Protein Data Bank (PDB, [40]) with the IDs listed in Table 1, which indicates the correspondence between UNIPROT IDs [41] and PDB IDs for the crystallographic and NMR structures used in this study. Simulations were prepared with Schrodinger’s Maestro GUI made available as part of the Desmond [42] software package (which also ran MD simulations) used Na^+^ or Cl^−^ ions to achieve electrical neutrality and the TIP4PEW water model. In order to avoid artifacts due to truncation of the simulated constructs and facilitate parameterization in AMBER99SB, the terminal amino acid residues present in the coordinate sets obtained from the PDB were capped. Simulations ran for 36 ns (following default relaxation/minimization protocols), with snapshots recorded every 14.4 ps (2500 snapshots). Re-parameterization of each simulation to use the AMBER99SB force field was performed using Desmond’s viparr utility. Simulations were run at room temperature (defined for each protein by the temperature at which NMR experiments used to solve the protein’s structure were performed: for all proteins in this study that temperature was very nearly 300K) in order to mimic the conditions in both the NMR tube and during (room temperature) crystallization as well as at 100K to mimic conditions obtained during cryo-cooled x-ray diffraction experiments. Dangling ends of proteins chains not present in the crystallographic coordinates deposited in the PDB were *not* filled in computationally but rather were omitted from each simulation.

**Table 1:** UniProt, NESG and PDB IDs of Studied Proteins and Protein Structures

| *ID #* |  | *PDB ID* |  |
| --- | --- | --- | --- |
| <u>UNIPROT</u> | <u>NESG</u> | <u>Crystal</u> | <u>NMR</u> |
| Q8ZRJ2 | StR65 | 2ES9 | 2JN8 |
| P74795 | SgR42 | 3C4S | 2JZ2 |
| Q7VV99 | BeR31 | 3CPK | 2K2E |
| Q8KFZ1 | CtR107 | 3E0H | 2KCU |

Initial parsing and visualization of each trajectory were performed using VMD [43]. Reformatting was completed for the multi-structure PDB file output from VMD into a multi-model format suitable for further analysis. THESEUS [29] superimposed MD trajectories prior to coordinate variance calculation and the MATLAB [44] implementation of the FindCore Toolbox superimposed NMR ensembles. MATLAB scripts also tabulated B-factor and coordinate variance/uncertainty data for analysis via Friedman’s test. Friedman’s test [45], also performed in MATLAB, is a non-parametric analog of ANOVA with repeated measures used here to compare whether coordinate uncertainties, variances and B-factors are significantly different for different atom types.

## Results and discussion

Fig 1 illustrates how coordinate uncertainty in NMR-derived “ensembles” (panel B) tracks coordinate variance in MD simulations (e.g. at 300K in panel D). The position of the carbonyl oxygen atom in residue 42 varies both across structural models in the NMR ensemble and over MD trajectories (panels C and D), and this oxygen atom is splayed more than the carbonyl carbon to which it is attached in panels B – D. However, the crystallographic B-factor for this carbonyl oxygen (22.15) is not particularly high nor is it much larger than that of the carbonyl carbon (21.83). Meanwhile on the opposite side of that peptide bond’s plane, the amide nitrogen from residue 43 is relatively well superimposed in the NMR ensemble and MD trajectory: the motion of the peptide plane appears to pivot around the amide nitrogen and proton. However, in the crystallographic structure, the B-factor (21.47) is barely lower than that of the carbonyl atoms.

**Fig 1:**
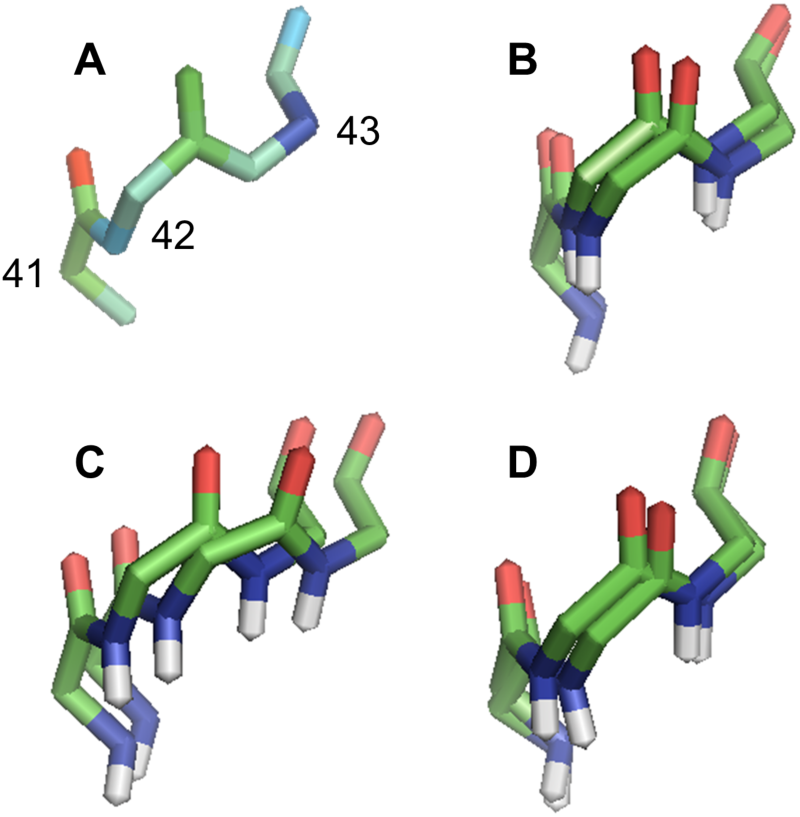
Backbone Traces of Residues 41–43 from Q8ZRJ2. (A) Crystallographic structure (PDB ID 2ES9) colored by B-factor with blue being low and red being high. Residue numbers shown in this panel reflect residue numbers in all panels. (B) FindCore superimposition of NMR ensemble (PDB ID 2JN8); this superimposition was calculated using a core atom set drawn from all heavy atoms (using all deposited models in the FindCore calculation) and not merely the residues shown. THESEUS superimposition, calculated from the entire MD trajectory using all heavy atoms, of MD trajectories simulated using the AMBER force field, showing snapshots 100 and 1000, at (C) 100K and (D) 300K. In panels (B) – (D), carbonyl oxygens are red, amide nitrogens are blue, carbons are green and amide hydrogens are white. Note the splaying in the carbonyl oxygens in panels (B) – (D) and the relatively well superimposed amide nitrogens in panels (B) and (D). Even in panel (C), amide nitrogens are better superimposed than carbonyl oxygens. In general, peptide planes appear to pivot with the amide protons and/or amide nitrogens being relatively imobile with the carbonyl oxygens at the opposite end of the peptide plane being relatively mobile. This pattern is not apparent in the B-factors depicted in panel (A).

Application of Friedman’s test to coordinate uncertainties, ranked from lowest to highest on a per-residue basis, of NMR structures confirms observations made in the development of Expanded FindCore [22] and those illustrated in Fig 1: in each of the four NMR ensembles analyzed (Fig 3, column A), carbonyl oxygen atoms are significantly likely to rank higher in coordinate uncertainty or variance than carbonyl carbons or amide nitrogens. Ranks of α-carbons also tended to be higher than, albeit not always significantly so, those of carbonyl carbons and amide nitrogens. This pattern is not an artifact of the FindCore method, but is also found following superimposition of NMR-derived ensembles with THESEUS [29] (Fig 2). No corresponding pattern exists with B-factors (Fig 3, column B), however: the only significant difference between B factor ranks was between the amide nitrogens and α-carbons of P74795.

**Fig 2:**
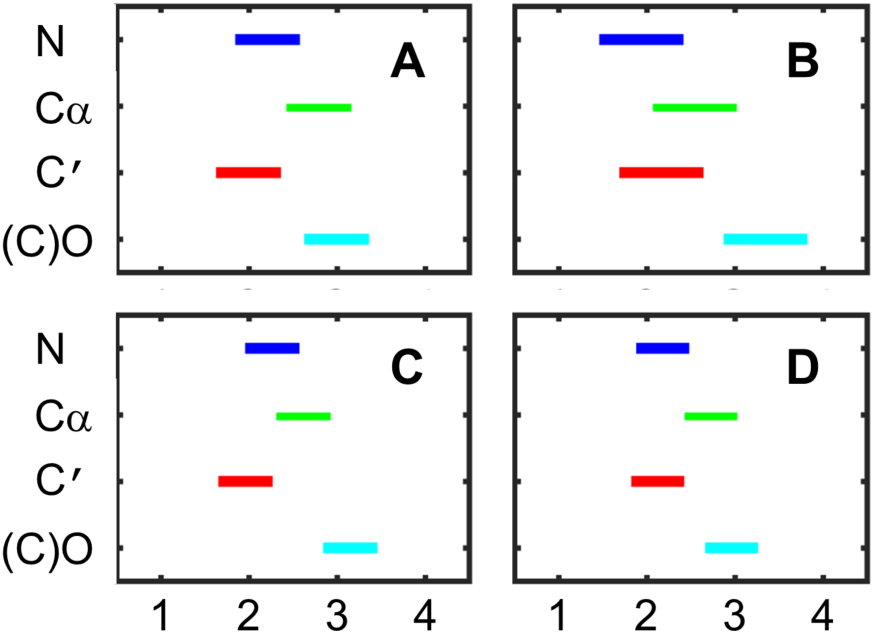
THESEUS Superimposed NMR Ensembles. This figure depicts the results of Friedman’s test applied to coordinate uncertainties for NMR structures as superimposed using THESEUS [28,29]. Panel (A) shows results from the superimposition of Q8ZRJ2 (PDB ID 2JN8), panel (B) for P74795 (PDB ID 2JZ2), panel (C) for Q7VV99 (PDB ID 2K2E), and panel (D) for Q8KFZ1 (PDB ID 2KCU). Note that as with the FindCore [25] superimpositions shown in Figure 3, carbonyl oxygen coordinates were significantly likely to be more uncertain than those for either the amide nitrogens or carbonyl carbons.

**Fig 3:**
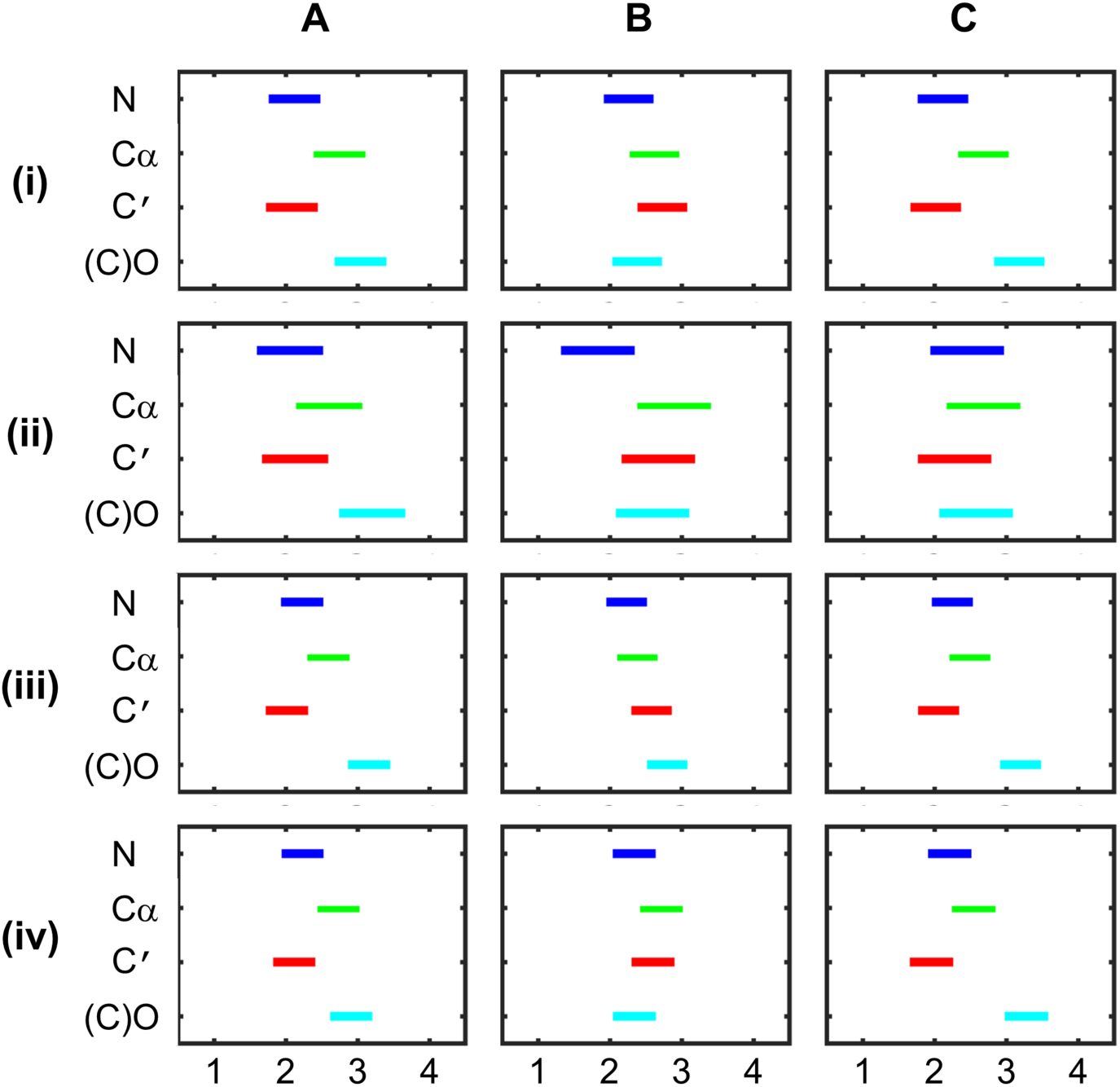
Comparison of Coordinate Uncertainties and B-factors for Four Proteins. This figure depicts the results of Friedman’s test applied to (Column A) coordinate uncertainties for NMR structures as superimposed using FindCore [26], (Column B) crystallographic B-factors, and (Column C) coordinate variances across a THESEUS [29,30] superimposed MD trajectory, this time seeded from the crystallographic structure but with selenomethionines replaced by methionines and simulated at 300K using the AMBER force field. Row (i) compares coordinate uncertainties, variances and B-factors for Q8ZRJ2, row (ii) is the comparison for P74795, row (iii) is for Q7VV99 and row (iv) is for Q8KFZ1. The bars in all panels depict the mean rank ± 3σ (x axes) for (y axes, from top to bottom in each panel) amide N, Cα, carbonyl C and carbonyl O atoms. Note that these are not true simultaneous confidence intervals, but non-overlapping intervals do indicate significant differences in mean rank (Z = 3, p = 0.13%). In all superimposed NMR ensembles and all superimposed MD trajectories except for the 300K AMBER trajectory for P74795, carbonyl oxygen coordinates were significantly likely to be more uncertain than those for either the amide nitrogens or carbonyl carbons.

In MD simulations ran at 300K and using AMBER force field (Fig 3, column C), carbonyl oxygens have a significant tendency to rank higher in coordinate variance than amide nitrogens and carbonyl carbons. Coordinate variances of α-carbons also tended to rank higher than coordinate variances for amide nitrogens and carbonyl carbons, but this effect was neither as strong nor as significant. This pattern exists in simulations using the AMBER force field at both 100K (Fig 4A) and 300K (Fig 4B) and in simulations using OPLS (at 100K) with selenomethionines maintained (Fig 4C) or replaced by methionines (Fig 4D). The persistence of this pattern across multiple MD simulations indicates that its absence in crystallographic B-factors is not due to the presence of selenomethionine residues in the constructs used in crystallography nor is its absence in crystallographic results due to a temperature effect, e.g. associated with cryogenic cooling during X-ray diffraction experiments.

**Fig 4:**
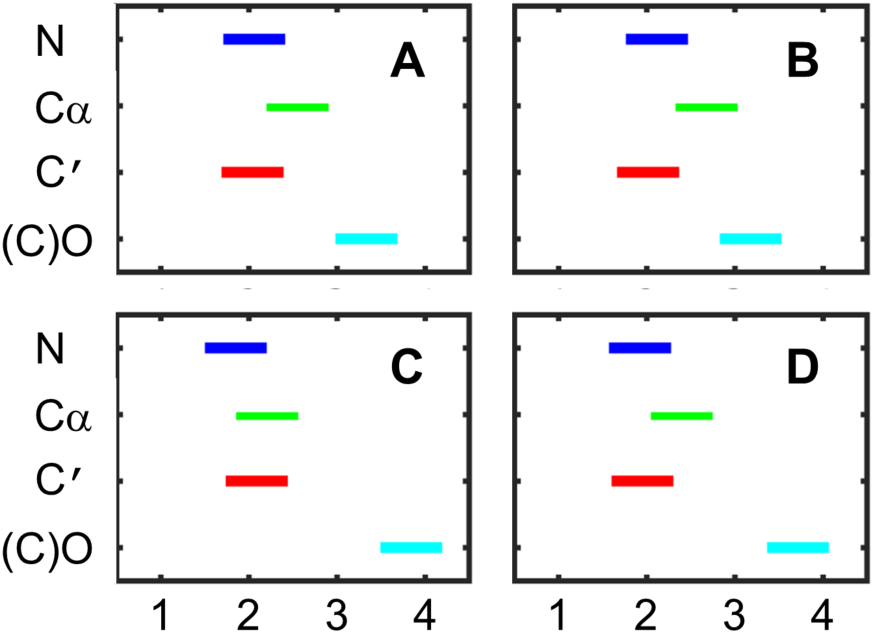
Comparison of MD Trajectories Seeded with Q8ZRJ2. This figure depicts the results of Friedman’s test applied to MD trajectories seeded with the protein uniprot ID: Q8ZRJ2 using the AMBER force field at 100K (panel A), the AMBER force field at 300K (panel B), the OPLS force field at 300K with selenomethionines (panel C), and the OPLS force field at 300K with selenomethionines replaced with methionine (panel D).

That carbonyl oxygens possess a significant tendency to have higher coordinate variances in THESEUS superimposed MD ensembles as well as having higher coordinate uncertainties across FindCore superimposed NMR “ensembles” indicates the pattern of coordinate uncertainties observed in NMR-derived structures is not solely an artifact of the superimposition method (THESEUS vs. FindCore), particular force field used (AMBER and OPLS in MD simulations, CNS [38,39] in NMR refinement) nor the particular characteristics of NMR-based structure determination (e.g. a lack of experimentally derived restraints on carbonyl oxygen atoms). The persistence of the tendency for carbonyl oxygens to have higher coordinate variability between ensembles explored via MD simulation and NMR-derived “ensembles”, which typically consist of models resulting from replicated simulated annealing calculations, indicates that this tendency is not solely an artifact of the structure sampling scheme used in NMR calculations.

Carbonyl oxygen uncertainties, as compared with those of carbonyl carbons and amide nitrogens, on average rank even higher in NMR derived structures than coordinate variances for carbonyl oxygens rank in MD trajectories. This indicates that NMR-based structure determination insufficiently restrains the positions of carbonyl oxygen atoms: perhaps NMR-based structure determination would benefit from additional restraints on the position of carbonyl oxygens, such as increasing the weight of hydrogen bonding restraints or taking into account other non-covalent interactions involving carbonyl oxygens such as those with aromatic rings [46]. Since even in MD simulations, carbonyl oxygens are relatively unrestrained, perhaps even MD simulations could benefit from better representation of hydrogen bonding or other non-covalent interactions, such as n➔π^*^ interactions [47], in MD force fields.

In addition to potentially inadequately representing quantum mechanical phenomena such as hydrogen bonding and n➔π^*^ interactions, many force fields strongly penalize any deviation of a peptide bond from planarity. In particular, requiring peptide bonds to remain planar may cause more complex motions of the amide backbone to be represented by simple rocking motions along an axis near the N-C bond axis but angled slightly toward the Cα. This motional model, by placing carbonyl oxygens furthest from the axis of motion (and Cα atoms second furthest), inappropriately represents them as being most mobile. Deficiencies in representing hydrogen bonding in force fields [48] may also be problematic when such deficiencies result in insufficient restraints on carbonyl oxygen positions; hydrogen bonds being important in stabilizing protein tertiary structure [49], may represent important restraints in carbonyl oxygen position across MD trajectories just as they are in NMR-based structure determination.

While NMR replicates the pattern of coordinate variances (variances tending to be highest in carbonyl oxygens) found in MD ensembles, this pattern is not found in Crystallographic B-factors. However, it may still be the case that overall, Crystallographic B-factors, which ideally arise from static and dynamic disorder, better reflect protein flexibility as calculated using MD simulations than do coordinate uncertainties calculated across ensembles of NMR-derived structures. After all, coordinate uncertainties calculated from FindCore superimpositions are statistical quantities measuring the precision of the NMR-derived coordinates rather than physical quantities directly attributable to protein dynamics. To remove the effects of the pattern in coordinate uncertainties/variances found in MD and NMR results, Fig 5 and Table 2 compare coordinate uncertainties, coordinate variances and Crystallographic B-factors only for Cα atoms. Using a single backbone atom type, Cα being a standard choice of backbone atom to use, rather than an extended set of backbone and/or side-chain atoms, obviates any correlations between coordinate uncertainties and coordinate variances due solely to the kinds of patterns analyzed above.

**Table 2:** Correlation Coefficients between Backbone Heavy Atom Coordinate Uncertainties (NMR), B-factors (Xtal) and Coordinate Variances from Superimposed MD Trajectories

| Protein | NMR v. | Xtal v. | AMBER 100K | OPLS MSE 100K |
| --- | --- | --- | --- | --- |
|  | AMBER 300K | AMBER 300K |  |  |
| Q8ZRJ2 | 0.6785 | 0.0137 | 0.097 | 0.1313 |
| P74795 | 0.8494 | 0.5892 | 0.6551 | 0.361 |
| Q7W7N7 | 0.7934 | 0.472 | 0.6071 | 0.5038 |
| Q8KFZ1 | 0.7672 | 0.7279 | 0.7607 | 0.1562 |

**Fig 5:**
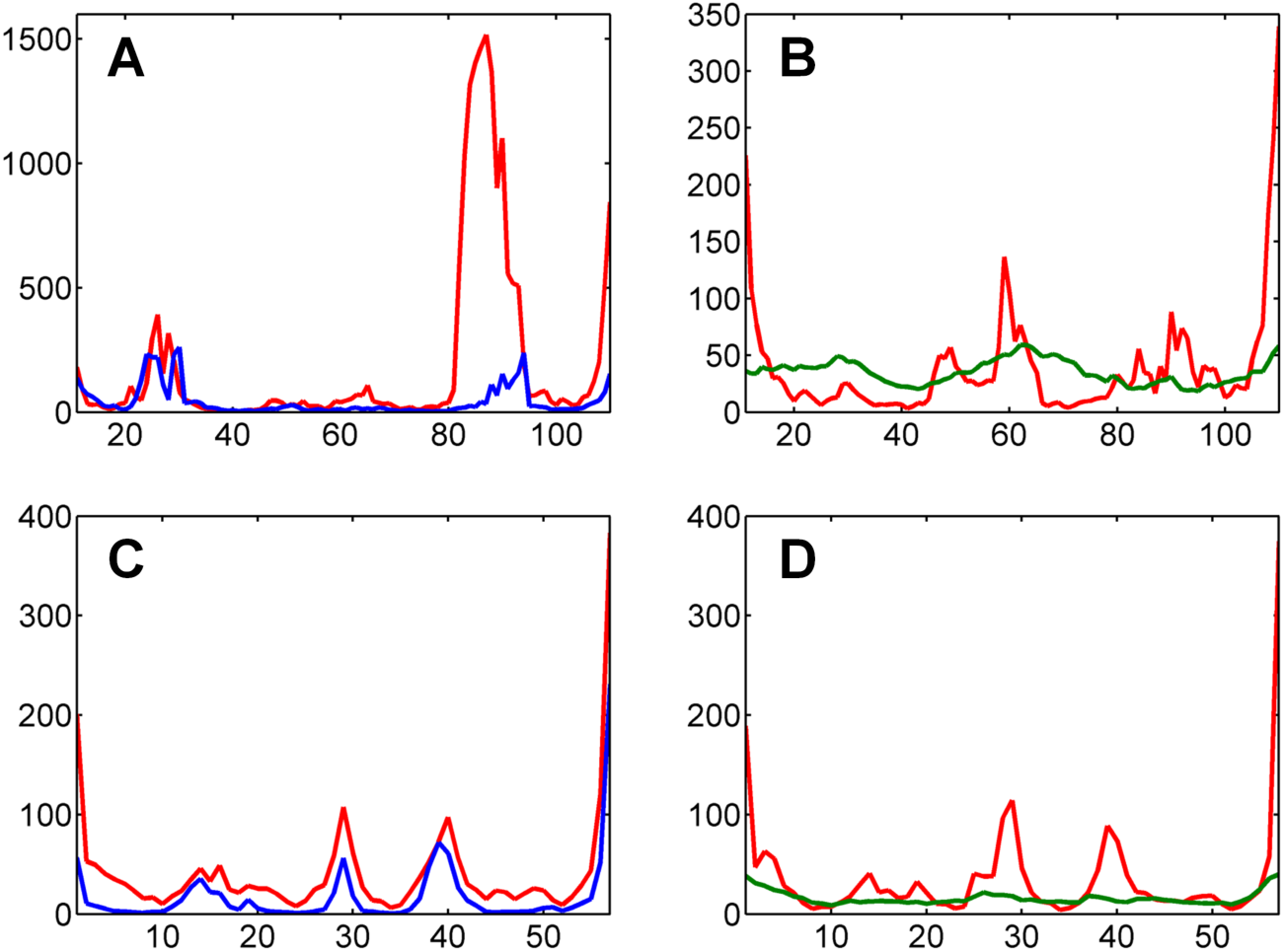
Coordinate variances, uncertainties and B-factors for Cα atoms. Coordinate variances across MD trajectories superimposed using THESEUS[28,29] (red), NMR ensembles superimposed using FindCore [26] (blue) and crystallographic B-factors (green). Multiplication by 8π^2^/3 gives coordinate variances and uncertainties the same scale as B-factors. (A) Coordinate variances from an MD trajectory for Q8ZRJ2 at 300K using the AMBER99SB force field compared with coordinate uncertainties from the NMR structure (PDB ID 2JN8). (B) Coordinate variances from an MD trajectory for Q8ZJR2 at 100K using AMBER99SB compared with B-factors (from PDB ID 2ES9). (C) Coordinate variances from an MD trajectory for P74795 at 300K using the AMBER99SB force field compared with coordinate uncertainties from the NMR structure (PDB ID 2JZ2). (D) Coordinate variances from an MD trajectory for P74795 at 100K using AMBER99SB compared with B-factors (from PDB ID 3C4S).

As shown in Fig 5, Cα coordinate uncertainties in NMR structure determination track MD coordinate variances better than B-factors do. As previously demonstrated [15], B-factor profiles generally agree with MD coordinate variances but are smoother and rarely reach the same high values found in MD coordinate variances. On the other hand, NMR coordinate uncertainty profiles have the same “spikiness” as MD coordinate variances even though NMR coordinate uncertainties still tend to underestimate the magnitude of fluctuations which occur in flexible regions of proteins in MD simulations.

The correlation coefficients in Table 2 quantify the tracking between NMR coordinate uncertainties and MD coordinate variances and the lack of similar correlation with B-factors. Subjecting the correlation coefficients in Table 2 to Friedman’s test finds a significant (p = 0.026) difference between the columns: in particular coordinate variances in MD simulations at 300K using the AMBER99SB force field correlate significantly better to NMR coordinate uncertainties than to crystallographic B-factors. Parametric analysis (paired t-test) detects another significant difference between the correlation coefficients in Table 2: the correlations between coordinate variances in MD simulations at 300K using the AMBER99SB force field and NMR coordinate uncertainties are significantly higher than those comparing MD simulations at 100K using the OPLS force field (with selenomethionine residues maintained in the crystallographic structures used to seed these calculations) and Crystallographic B-factors. While the correlation coefficients between Crystallographic B-factors and coordinate variances in the MD simulations at 100K using the AMBER99SB force field are not significantly worse than the correlations between coordinate variances in AMBER99SB MD simulations at 300K and coordinate uncertainties across NMR structures, the correlation coefficients between MD and NMR are higher than the correlation coefficients between the 100K AMBER99SB coordinate variances and B-factors. In order words, coordinate uncertainties across NMR-derived structural ensembles reflect actual protein flexibility, as predicted using MD simulations, at least as well as Crystallographic B-factors do.

Similar to the case of the pattern in backbone heavy atom coordinate variances/uncertainties, the better correlation of protein flexibility simulated in MD calculations to NMR-derived coordinate uncertainties than to B-factors may reflect deficiencies common to the force fields used to calculate NMR-derived structures and MD trajectories. The high quality of crystallographic structures and the sheer number of “reflections” used to calculate crystallographic structures ideally mean that Crystallographic B-factors, while reflecting model quality which is in turn dependent on the quality of the force fields used in structure refinement, are less force field dependent than MD simulations and NMR-based structure calculations. Thus it may be that Crystallographic B-factors better describe protein flexibility than do MD calculations.

On the other hand, even in an ideal case where Crystallographic B-factors arise entirely from static and dynamic disorder, these B-factors reflect protein dynamics in the crystalline state and not in the solution state [50]. For example, previous studies comparing crystallographic B-factors and protein flexibility in solution indicate that crystallization has a “flattening” effect on protein flexibility [51]. The B-factor profiles shown in Fig 5 similarly display a smoother profile than either coordinate variances in superimposed MD trajectories or coordinate uncertainties from superimposed NMR “ensembles”. Moreover, since Debye-Waller theory attributes any reduction in diffraction pattern intensities relative to those expected given a static protein structure to local, harmonic motion, other processes that reduce diffraction pattern intensities may result in over-or even under-estimation of protein flexibility [52]. Relatedly, values obtained for B-factors are dependent on the refinement techniques used in interpreting X-ray data [51].

Nevertheless, the patterns described in this paper as well as the relatively high correlations between the *statistical* coordinate uncertainties derived from NMR and the putatively *physical* coordinate variances across MD ensembles may very well indicate deficiencies common to all force fields. Fully exploring the pervasiveness of the patterns described in this paper necessitates MD simulations and analysis of NMR structures beyond the four systems studied here. However, the analysis presented in this paper identifies that coordinate variances/uncertainties from at least some MD trajectories and NMR ensembles have properties not found in B-factors. This divergence between B-factors and coordinate variances potentially indicates that there remain critical concerns in force field development. Future studies of MD trajectories will hopefully reveal which potentially inaccurate aspects of force fields, such as the requirement that peptide bonds remain planar and inadequacies in the representation of non-covalent interactions such as hydrogen bonding as well as solvent/protein interactions, need the most adjustment. Addressing such deficiencies in force field construction can result in better descriptions of protein structure and hence facilitate the accurate prediction of protein dynamics, structure and folding pathways.

## Acknowledgements

The authors graciously acknowledge Adrian Roitberg for his insights into the results presented by DAS at the ACS Spring 2014 Meeting in Dallas. The authors also thank Gaetano Montelione for his insights into NMR-based structure determination as well as for his helpful comments and constructive discussion. D.A.S. thanks John Chodera for suggesting submission of this manuscript to bioRXiv. An award of Assigned Release Time for research from the Office of the Provost of William Paterson University of NJ facilitated completion of this work.

